# Neuronal differentiation through GPI-anchor cleavage by GDE2: regulation via sequence-dependent trafficking

**DOI:** 10.1101/532499

**Authors:** Fernando Salgado-Polo, Michiel van Veen, Bram van den Broek, Daniela Leyton-Puig, Anastassis Perrakis, Wouter H. Moolenaar, Elisa Matas-Rico

## Abstract

GDE2 (GDPD5) is a multispanning membrane glycerophosphodiesterase that cleaves select glycosylphosphatidylinositol (GPI)-anchored proteins and thereby alters cell behavior. GDE2 promotes neuronal differentiation and survival in cell-autonomous and non-cell-autonomous manners through cleavage of glypicans and RECK, respectively. Moreover, GDE2 is a prognostic marker in neuroblastoma and protects against progressive neurodegeneration in mice. However, its regulation remains unclear. Here we report that in neuronal cells, GDE2 undergoes constitutive endocytosis and travels back along both fast and slow recycling routes. This trafficking is differentially regulated by distinctive C-terminal sequences that modulate GDE2’s ability to release glypican-6 and induce neuronal differentiation. Specifically, we define a GDE2 truncation mutant that displays aberrant recycling and is dysfunctional, whereas a consecutive 10-amino acid deletion results in cell-surface retention and gain of function. These results highlight the importance of sequence-dependent membrane trafficking for GDE2 neuronal function, of relevance for nervous system disorders where trafficking has gone awry.

## Introduction

The surface of eukaryotic cells contains a great variety of glycosylphosphatidylinositol (GPI)-anchored proteins that are involved in the regulation of vital cellular functions, including signal transduction, cell adhesion, intercellular communication and differentiation. GPI-anchoring is a highly complex post-translational modification that tethers membrane proteins via their C-terminus to a unique glycosylated phosphatidylinositol (PI) core in the outer leaflet of the plasma membrane, particularly at lipid raft nanodomains (Ferguson et al., 2015; Fujita and Kinoshita, 2010; Paulick and Bertozzi, 2008). Since they lack a transmembrane domain, GPI-anchored proteins cannot transmit signals by themselves but must interact with transmembrane effectors or cellular adhesion pathways to achieve signaling competence. Importantly, GPI-anchoring confers a unique property to membrane proteins, namely susceptibility to phospholipase attack. Indeed, GPI-anchored proteins can be released from their anchor and detected as soluble forms in culture media and body fluids. Yet, identification of the responsible phospholipase(s) and their biological function has long been elusive.

Recent studies have advanced the field by showing that members of the glycerophosphodiester phosphodiesterase (GDPD) family, notably GDE2 and GDE3, function as GPI-specific phospholipases that cleave select GPI-anchored proteins (*in cis*) and thereby alter biological signaling cascades and cellular phenotype (Matas-Rico et al., 2016; Park et al., 2013; van Veen et al., 2017). GDE2 (also known as GDPD5) is the best studied family member and, along with GDE3 and GDE6, is characterized by six transmembrane helices, a catalytic GDPD ectodomain and intracellular N- and C-terminal tails (**Fig. 1A,B**). GDE2 acts in a phospholipase D (PLD)-like manner towards soluble substrates, *i.e.* releasing choline from glycerol-3-phosphocholine (Gallazzini et al., 2008), in common with virtually all GDPD family members (Corda et al., 2014; Ohshima et al., 2015). One notable exception is GDE3, which functions as a phospholipase C (PLC) and shows different substrate selectivity from GDE2 (Corda et al., 2009; van Veen et al., 2017).

**Figure 1.**
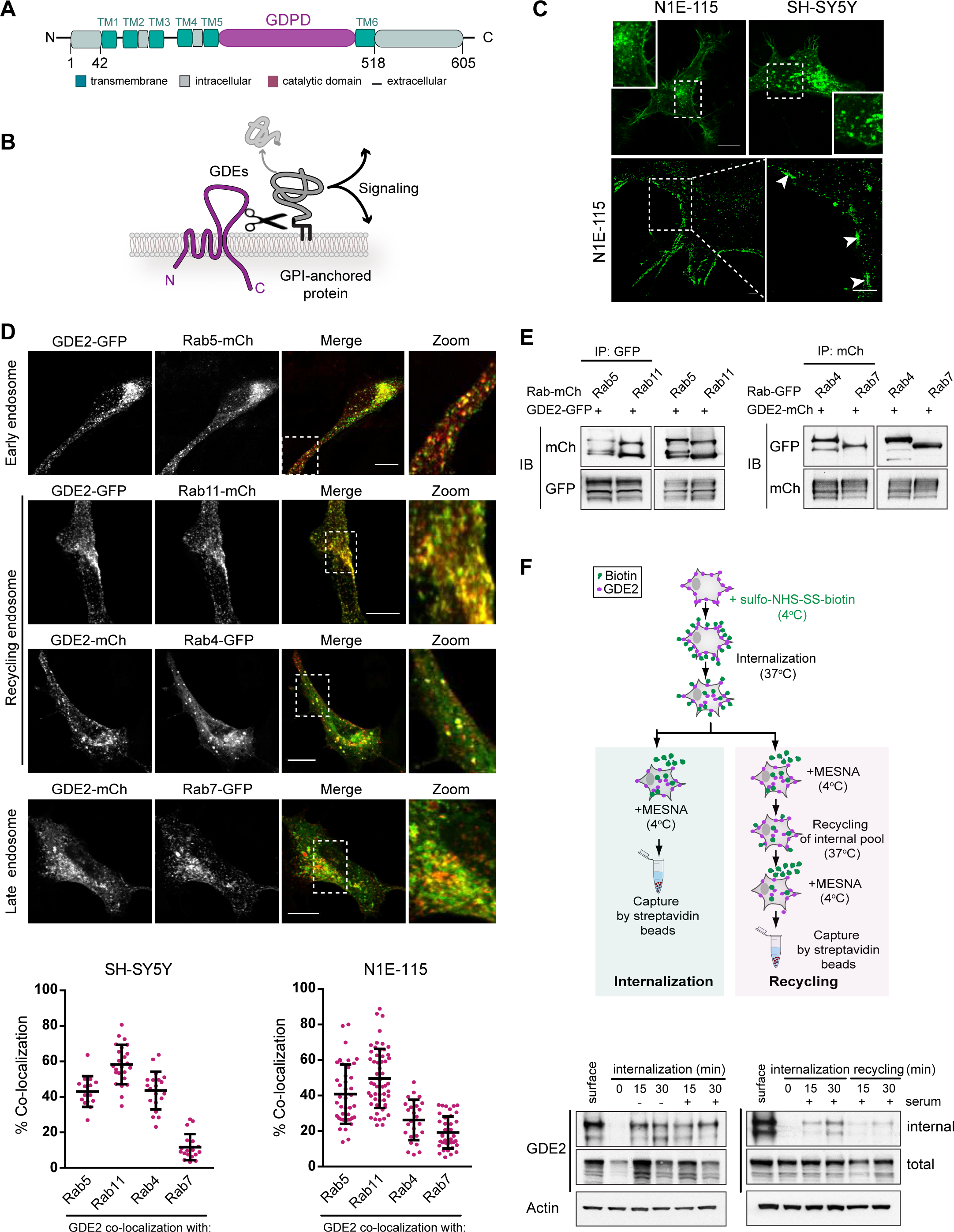
GDE2 localization and endocytic trafficking routes. **A.** Domain structure of GDE2 showing six transmembrane (TM) domains, a GDPD ectodomain and intracellular N- and C-terminal tails. **B.** GDE2 cleaves and sheds GPI-anchored proteins resulting in activation of signaling cascades. **C.** GDE2 subcellular localization. <u>Top</u>: Confocal images showing GDE2-GFP in membrane microdomains and intracellular vesicles in N1E-115 and SH-SY5Y cells; bar, 10 μm. <u>Bottom</u>: Super-resolution images of N1E-115 cells expressing GDE2-GFP. White arrows point to membrane microdomains. Bar, 1 μm. See also (Matas-Rico et al., 2016). **D.** Confocal images of GDE2 in early, recycling and late endosomes, Rab5-, Rab11-, Rab4-, and Rab7-positive, respectively, in SH-SY5Y cells. Bar, 10 μm. <u>Bottom panels</u> show quantification of GDE2 co-localization with the indicated Rab GTPases, expressed as the percentage of yellow versus red pixels (≥25 cells from three independent experiments). Data represent the median ± S.D. (error bars) of co-localization. **E.** GDE2 (fused to GFP or mCh) associates with the indicated Rab GTPases in HEK293T cells. GDE2 was immunoprecipitated (IP) and subjected to immunoblotting (IB) using either anti-GFP or anti-mCh antibody. **F.** <u>Top panel:</u> Schematic illustration of the internalization and recycling assay using biotin labelling. N1E-115 cells expressing GDE2-mCh were surface-labelled with NHS-S-S-Biotin. Internalization proceeded for 15 and 30 min at 37°C in presence or absence of 10% FBS. Surface biotin was reduced with MesNa at 4°C, and the cells were shifted to 37°C for the indicated time periods to trigger recycling of the internal pool. <u>Bottom panel</u>: The amount of internalized and total biotin-labelled GDE2 determined by immuno-blotting using anti-mCh antibody. Actin was used as loading control.

GDE2 was originally shown to drive neuronal maturation and survival in the developing spinal cord (Rao and Sockanathan, 2005; Sabharwal et al., 2011). Thereafter, GDE2 was found to mediate cleavage of GPI-anchored RECK, an ADAM10 metalloprotease inhibitor and Notch ligand regulator, leading to inhibition of Notch signaling and induction of differentiation in adjacent neural progenitors (Park et al., 2013). More recently, we reported that GDE2 promotes neuronal differentiation in a cell-autonomous manner through cleavage of glypican-6 (GPC6) (Matas-Rico et al., 2016). Glypicans are GPI-anchored heparan sulfate proteoglycans that play key roles in morphogenesis and can signal via growth factor recruitment or binding to receptor protein tyrosine phosphatases (Capurro et al., 2017; Farhy-Tselnicker et al., 2017; Ko et al., 2015). Enforced GDE2 expression led to altered Rho GTPase signaling, upregulation of neural differentiation markers, neurite outgrowth and resistance to neurite retraction; furthermore, GDE2 expression was found to strongly correlate with clinical outcome in neuroblastoma, a neurodevelopmental malignancy characterized by impaired differentiation (Matas-Rico et al., 2016). Furthermore, importantly, *Gde2* knockout mice display progressive neurodegeneration in the spinal cord with pathologies reflecting human neurodegenerative disease, including behavioral motor deficits, which was accompanied by reduced glypican release (Cave et al., 2017). Finally, depletion of GDE2 in zebrafish embryos causes motility defects and reduced pancreas differentiation, which was tentatively attributed in a part to enhanced Notch signaling; overexpression of human GDE2, but not catalytically dead GDE2, similarly led to developmental defects (van Veen et al., 2018). Taken together, these results underscore the need for tight control of GDE2 surface expression and activity *in vivo*. However, it is still unknown how GDE2 is regulated.

Here we report that in neuronal cells, GDE2 is regulated by membrane trafficking under tight control of its C-terminal tail. Membrane trafficking is a crucial regulatory process gone awry in neurodegenerative diseases. We identify unique C-terminal tail sequences that govern GDE2 export, endocytosis and recycling preference, respectively, and thereby modulate GDE2 activity both positively and negatively, as measured by GPC6 release and induction of neuronal differentiation marker genes. Our results reveal a mechanistic link between sequence-dependent membrane trafficking and GDE2 neuronal function, providing new insight into the control of GDE2-mediated GPI-anchor hydrolysis and its biological outcome with potential pathophysiological implications.

## Results

### GDE2 trafficking: constitutive endocytosis and recycling

We previously showed that in undifferentiated neuronal cells, GDE2 is enriched in recycling endosomes and co-traffics with the endogenous transferrin receptor, a prototypic cargo of clathrin/dynamin-dependent recycling, suggesting regulation through endosomal trafficking (Matas-Rico et al., 2016). We analyzed the molecular determinants of GDE2 trafficking using SH-SY5Y and N1E-115 neuronal cells, which express very low levels of endogenous GDE2. At the plasma membrane, GDE2 (HA-, GFP- or mCherry-tagged) is detected in discrete microdomains or clustered nanodomains, particularly in developing neurite tips, as revealed by super-resolution microscopy (Matas-Rico et al., 2016) (**Fig. 1C)**. Intracellularly, GDE2 is abundantly present in vesicles, particularly in the perinuclear region (**Fig. 1C).** Treatment with the dynamin inhibitor dynasore resulted in GDE2 accumulation at the plasma membrane with almost complete loss of GDE2-positive vesicles, confirming that GDE2 undergoes dynamin-dependent internalization instead of bulk endocytosis (**Fig. S1A**). GDE2-GFP-containing vesicles are highly mobile and show rapid directional movement towards the tips of developing neurites or micro-spikes, as shown by live-imaging (**Movie S1**).

Rab GTPases are master regulators of membrane trafficking and play key roles in maintaining neuronal function (Mignogna and D’Adamo, 2018; Wandinger-Ness and Zerial, 2014). In both neuronal cell lines, GDE2-GFP co-localized with early endosome marker Rab5-mCh (**Fig.1D** and **Fig. S1B**). But the majority of intracellular GDE2 was detected in two distinct populations of recycling endosomes, namely those representing the Rab4- and Rab11-regulated recycling routes, “fast” and “slow**”**, respectively (**Fig. 1D and S1B**). Rab4-mediated fast recycling of membrane cargo involves a half-time of a few minutes, whereas Rab11 regulates slow recycling through perinuclear endosomes with a half-time of >10 min. (Maxfield and McGraw, 2004). A relatively small percentage of GDE2 was detected in Rab7-positive (Rab7^+^) late endosomes, or multivesicular bodies, which deliver cargo to lysosomes for proteolytic degradation. Consistent with this, a small fraction of GDE2 was found in LAMP1-positive lysosomes **(Fig. S2)**.

Quantification of the results from both neuronal cell lines showed that GDE2 predominantly localized to Rab11^+^ endosomes (mean ∼60%), somewhat less to Rab5^+^ and Rab4^+^ endosomes (mean 30-45%, depending on the cell line), and much less to Rab7^+^ late endosomes and lysosomes (**Fig. 1D** and **Fig. S2**). To validate the GDE2 localization data biochemically, we examined the interaction of GDE2 with relevant Rab GTPases in HEK293 cells. GDE2 immuno-precipitates blotted for Rab proteins showed GDE2 in complex with Rab4, Rab5, Rab7 and Rab11, in support of the co-localization results (**Fig. 1E)**. We next measured GDE2 internalization and recycling using a biotin labeling procedure (**Fig. 1F**). To this end, GDE2-mCh-expressing N1E-115 cells were surface-labeled with NHS-SS-Biotin at 4° C and endocytosis was initiated by a temperature shift to 37° C. As shown in **Fig. 1F**, internalized GDE2 was found to recycle back to the plasma membrane within 15-30 min. Of note, the GDE2 intracellular pool was not affected by serum stimulation (**Fig. 1F**). Similarly, GDE2 co-localization with Rab4, Rab5 and Rab11 was not altered in the presence of serum (results not shown). We therefore conclude that GDE2 endocytosis and ensuing recycling is a constitutive process, insensitive to serum factors.

### C-terminal tail truncations uncover unique regulatory sequences

Many integral membrane proteins contain linear sequence motifs that determine endocytosis, recycling or degradation (Bonifacino and Traub, 2003; Cullen and Steinberg, 2018). However, sequence inspection did not reveal canonical sorting motifs in the cytoplasmic regions of GDE2. To explore the sequence determinants of GDE2 trafficking, we focused on the 88-aa long C-terminal tail (aa 518-605) (**Fig. 2A**). Of note, the GDE2 distal C-terminal region (aa 570-605) shows marked sequence divergence among vertebrates (**Fig. 2A**), suggesting that the last not well-conserved C-terminal residues do not play a key regulatory role but could have a species-specific function.

**Figure 2.**
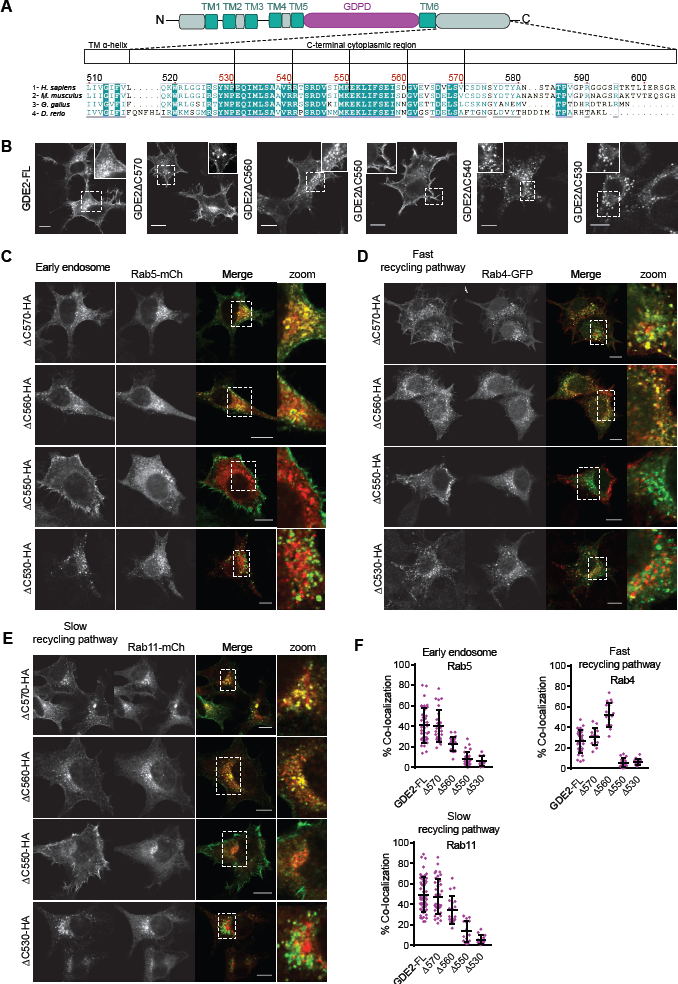
C-terminal tail truncations uncover unique regulatory sequences. **A.** C-terminal sequence alignments of human, mouse, chicken and zebrafish GDE2, and the truncations made in human GDE2. **B.** Subcellular localization of GDE2-HA and its truncation mutants in N1E-115 cells. Bar, 10 μm. **C**,**D**,**E**. Confocal images of N1E-115 cells co-expressing the indicated GDE2 CT truncations and Rab5-mCh (**C**), Rab4-GFP (**D**) and Rab11-mCh (**E**). Bar, 10 μm. **F.** Quantification of GDE2-Rab co-localization (percentage of yellow versus red pixels for ≥25 cells from three independent experiments). Data represent the median ± S.D. (error bars) of co-localization.

We made GDE2 C-terminal truncations at aa 570, 560, 550, 540 and 530 (HA-tagged), respectively (**Fig. 2A)**, expressed them in N1E-115 cells and analyzed their co-localization with Rab5, Rab4 and Rab11. GDE2(ΔC570) showed the same subcellular localization as full-length GDE2, indicating that the last 35 residues (aa 571-605) are dispensable for proper GDE2 localization and trafficking (**Fig. 2B**). GDE2(ΔC560) showed reduced surface expression (see also **Fig. 3A,B** below) and less Rab5 co-localization, but it accumulated preferentially in Rab4^+^ fast recycling endosomes at the expense of Rab11 co-localization (**Fig. 2B-F**). This suggests that the aa (561-570) region of GDE2 is required for endocytosis and, intriguingly, determines recycling pathway preference, redirecting endocytosed GDE2 from slow to fast recycling. Strikingly, a consecutive 10-aa truncation caused GDE2(ΔC550) to be retained at the plasma membrane with little or no protein detected in early and recycling endosomes (**Fig. 2C-E**). It thus appears that aa (551-560) region confers endocytosis competence and ensuing recycling to GDE2.

**Figure 3.**
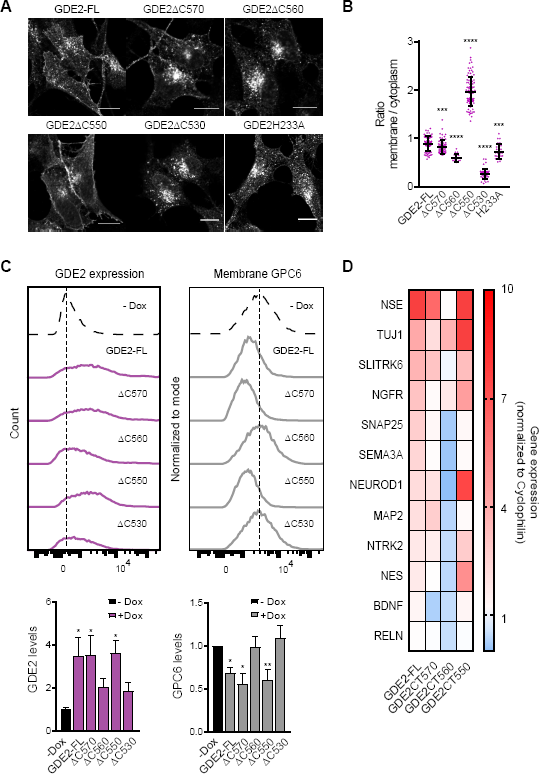
Sequence-dependent trafficking determines GDE2 activity. **A.** Localization of GDE2 and the indicated CT truncation construct in Dox-treated SH-SY5Y cells (48 hrs). **B.** Quantification of GDE2 surface versus cytosol localization in inducible SH-SY5Y cells (24 hrs). At least 20 cells from three independent preparations were segmented and analysed by IMAGEJ to calculate the membrane/cytoplasmic ratio (median ± SEM); ***p<0.001, ****p<0.0001, unpaired t test. **C.** <u>Left panel,</u> Surface expression of GDE2-HA and its truncated mutants upon Dox-induced (48 hrs) expression, detected by flow cytometry. <u>Right panel,</u> GPC6 surface levels as a function GDE2-HA expression compared to GDE2-deficient cells (-Dox), detected by flow cytometry using GPC6 antibody. In both cases, representative histograms from the same experiment are shown. <u>Lower panels</u>, Quantifications (mean ± SEM) of the above FACS data from three independent experiments; *p<0.05; **p<0.01, paired t test. **D.** Induction of neuronal differentiation marker genes upon Dox-induced expression of the indicated GDE2 constructs as determined by qPCR and shown in a heat map. For quantification see **Fig. S3C**.

Finally, upon further truncation at residue 540 or 530, GDE2 was no longer detected at the plasma membrane, but remained trapped intracellularly as aggregates, suggestive of faulty expression and misfolding (**Fig. 2B-F**). Misfolded proteins are usually routed to the autophagosome-lysosome degradation pathway; consistently, GDE2(**Δ**C530) was found to accumulate in lysosomes (**Fig. S2**). We therefore conclude that the aa 530-550 region is required for proper GDE2 expression, folding and transport through the early secretory pathway. The respective GDE2-Rab co-localization results are quantified in **Fig. 2F**. Together, these results uncover unique regulatory sequences in the C-terminal tail of GDE2.

### Sequence-dependent trafficking determines GDE2 activity

To determine how GDE2 trafficking and subcellular localization relates to activity, we used SH-SY5Y cells stably expressing doxycycline (Dox)-inducible GDE2 (Matas-Rico et al., 2016). GDE2-HA constructs were analyzed for their expression and subcellular localization upon Dox treatment. Full-length (FL) GDE2, its deletion mutants and catalytically dead GDE2(H233A) showed maximal expression after 24-48 hrs of Dox treatment; it is also seen that GDE2(**Δ**C540) and GDE2(**Δ**C530) were very poorly expressed (**Fig. S3A**), consistent with their exocytosis failure and apparent misfolding. After Dox treatment of SH-SY5Y cells, the GDE2 truncation mutants showed differential distribution between plasma membrane and endocytic vesicles very similar to that in N1E-115 cells (**Fig. 3A**). Catalytically dead GDE2(H233A) showed the same subcellular distribution as wild-type GDE2, which we take as evidence that GDE2 trafficking is independent of its GPI-hydrolyzing activity. We measured surface levels of GDE2 and its mutants relative to their intracellular accumulation by quantitative analysis of confocal images (**Fig. 3B**). GDE2-FL and GDE2(**Δ**C570) displayed similar surface to cytoplasm ratios, whereas GDE2(ΔC550) showed a much higher ratio, consistent with its preferential retention at the plasma membrane and failure to undergo endocytosis. GDE2(ΔC560) showed a reduced surface to cytoplasm ratio, in agreement with its preferred accumulation in recycling endosomes, particularly in the Rab4^+^ fast recycling route.

We then measured the surface expression of GDE2-HA and its truncation mutants, and their efficacy of inducing GPC6 release, using flow cytometry (**Fig. 3B,C)**; catalytically dead GDE2(H233A) was used as a negative control (**Fig. S3B**). GDE2-FL, GDE2(ΔC570) and GDE2(ΔC550) showed similar surface levels, while GDE2(ΔC560) showed reduced surface expression. Catalytic activity of GDE2 and its truncation mutants towards GPC6 correlated with their respective surface levels, except for GDE2(**Δ**C560) (**Fig. 3C).** Interestingly, the latter mutant was markedly inefficient in shedding GPC6 from the plasma membrane, correlating well with reduced surface expression levels and its preference for fast recycling (**Fig. 2F**). We conclude that the aa 551-560 region negatively regulates GDE2 function by conferring a recycling pathway imbalance and failure to attack GPC6.

GDE2 regulates a neuronal differentiation program as revealed by differential gene expression analysis (Matas-Rico et al., 2016). We investigated how the above GDE2 truncation mutants affected the expression of a set neuronal differentiation marker genes. These included the neurogenic transcription factor *NEUROD1*, neuron-specific enolase and cytoskeletal proteins (*NSE, NES, MAP2*), neurotrophic receptors (*NTRK2, SLITRK6*) and others. Induction of these genes by the respective GDE2 truncation mutants mirrored the induction of GPC6 release, albeit to varying degrees (**Fig. 3D** and **Fig. S3C**). GDE2(ΔC570) was fully active in most cases, similar to GDE2-FL. Strikingly, again, GDE2(ΔC560) was functionally inactive as it failed to induce significant gene transcription, correlating with its aberrant recycling behavior and lost ability to shed GPC6 (**Fig. 3D** and **Fig. S3C**). Finally, GDE2(ΔC550) showed gain of function towards most of the above differentiation markers, which we attribute to endocytosis failure and retention on the plasma membrane.

## Discussion

Membrane trafficking is of vital importance for cellular homeostasis and numerous signaling processes, particularly in the nervous system, and its dysregulation can lead to disease. Here, we have uncovered unique sequence determinants of GDE2 trafficking and their impact on GDE2 activity in undifferentiated neuronal cells. As summarized in **Fig. 4(A,B)**, full-length GDE2 and GDE2(ΔC570) show similar subcellular localization and travel along both fast and slow recycling routes, and have similar differentiation-inducing activity. It thus appears that the last 35 C-terminal residues (aa 571-605) are dispensable for GDE2 trafficking and function. Yet, this rather poorly conserved region could play a species-specific role. Indeed, in zebrafish, knockdown of C-terminally truncated GDE2 (lacking 34 residues), encoded by *Gdpd5b*, causes less severe developmental defects than observed with *Gdpd5a* knockdown (van Veen et al., 2018).

**Figure 4.**
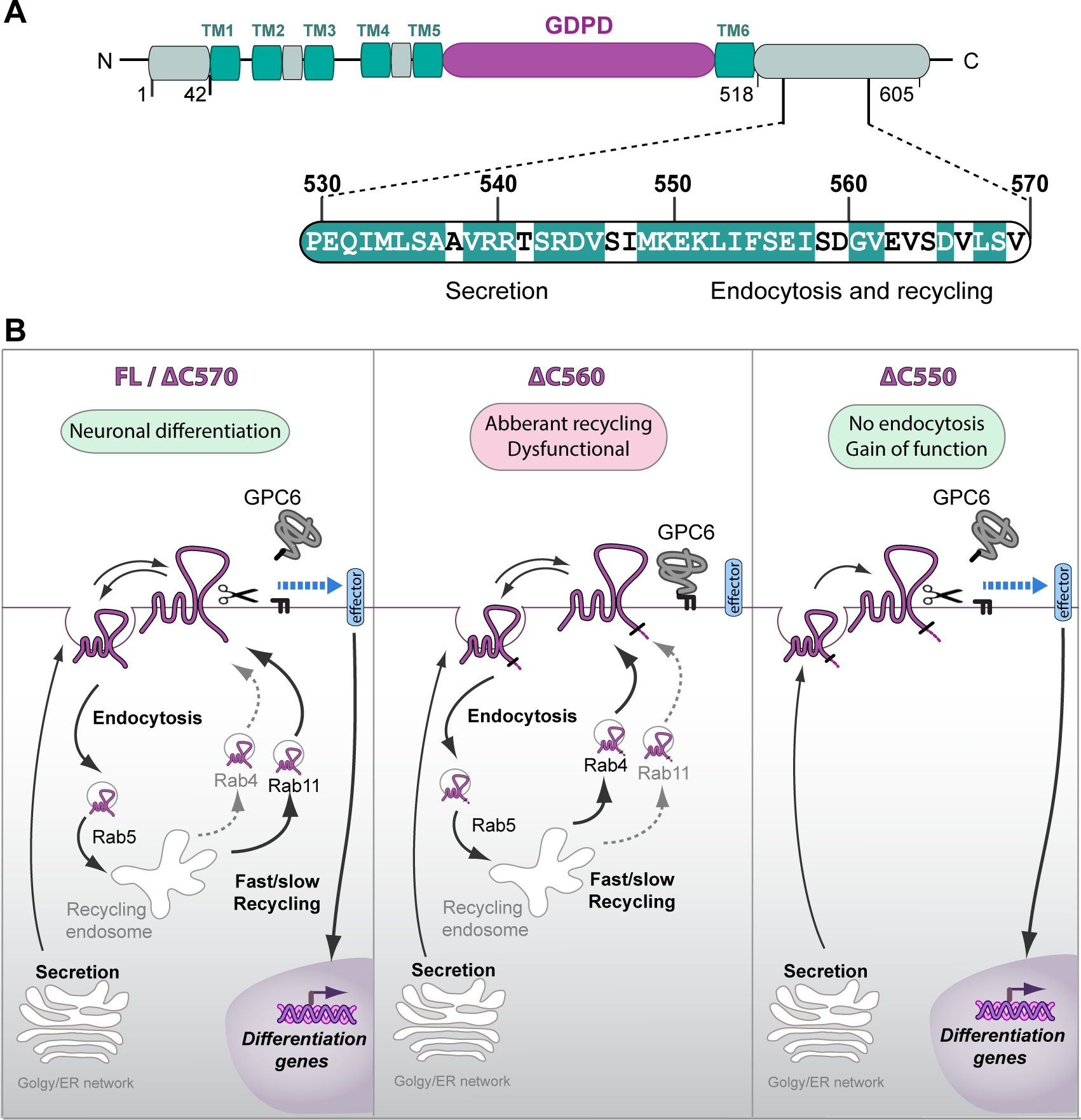
Sequence determinants of GDE2 trafficking and biological activity. **A.** The C-terminal region (aa 530/541-550) is essential for proper expression, secretion and membrane insertion. Sequence (aa 551-560) determines endocytosis and Rab4 fast recycling preference, but negatively regulates GDE2 function. Sequence (561-570) is required for proper GDE2 recycling and function. Residues in white are not conserved between mammalian, chicken and/or zebrafish GDE2 (cf. **Fig. 2A**). **B.** Scheme of membrane trafficking, localization and biological output of GDE2 and the indicated truncation mutants, acting in a cell-autonomous manner through GPC6 cleavage and an as-yet-unknown transmembrane effector. GDE2 is constitutively internalized while the majority of endocytosed GDE2 recycles along Rab4^+^ and Rab11^+^ routes in a sequence-dependent manner. Small part of GDE2 is sorted to Rab7^+^ late endosomes (not illustrated). GDE2(ΔC560) shows preference for Rab4-driven fast recycling and is dysfunctional, whereas GDE2(ΔC550) is retained at the cell surface with gain of function. Signaling efficacy is inferred from GPC6 shedding and induction of neuronal differentiation marker genes.

Upon further truncation, the resulting GDE2(ΔC560) mutant is redirected from Rab11^+^ slow to Rab4^+^ fast recycling and shows reduced surface expression but, intriguingly, is virtually inactive (**Fig. 4B**). Strikingly, deletion of the next 10 amino acids, generating GDE2(ΔC550), confers cell-surface retention attributable to endocytosis failure and a marked gain-of-function phenotype. This implies that the 551-560 sequence is essential for GDE2 exocytosis and plasma membrane insertion but lacks endocytosis information. Finally, juxta-membrane region (aa 518-550) is required for GDE2 biosynthesis and export, as the ΔC540 and ΔC530 mutants are detected as aggregates and fail to enter the secretory pathway.

The most striking regulatory sequence emerging from our analysis is 551-EKLIFSEISD-560, as it contains information that shifts the Rab11/Rab4 recycling balance and renders GDE2 dysfunctional, and therefore qualifies as a negative regulatory sequence determinant. How to explain that ΔC560 is inactive, whereas ΔC550 is biologically “hyperactive”? At this stage, we can only speculate about underlying mechanisms. Aberrant recycling may misdirect GDE2 to nanodomains that are short of GPC6 substrate. Alternatively, due to its fast recycling, GDE2(ΔC560) might have a relatively short residence time at the plasma membrane, thereby preventing effective GDE2-substrate interaction. Whether the latter mutants may undergo differential sorting to Rab7^+^ late endosomes remains to be determined, although it concerns a relatively small percentage of the total endosomal GDE2 pool. To gain further insight into the molecular determinants of GDE2 trafficking, it will be essential to identify adaptor proteins and effectors that interact with defined C-terminal sequence motifs to drive the endocytic sorting machinery. Post-translational modifications could also modulate GDE2 trafficking, as exemplified by the finding that under oxidative conditions, disulfide bonding disrupts GDE2 export to the plasma membrane (Yan et al., 2015). Aside from tightly regulated trafficking, the biological outcome of GDE2 activity will critically depend on the local availability of GPI-anchored substrates and their spatiotemporal organization in a given cell type. Determination of the substrate specificity of GDE2 and its structural basis has therefore high priority.

Since GDE2 deficiency leads to neurodegeneration in mice with pathologies analogous to human disease and, furthermore, is associated with poor outcome in neuroblastoma, GDE2 dysfunction might underlie aspects of neurodegenerative disease and/or the pathophysiology of neurodevelopmental disorders such as neuroblastoma. Yet, disease-associated GDE2 deficits have not been documented to date, although *GDE2* (*GDPD5*) expression is decreased in motor neurons from amyotrophic lateral sclerosis (ALS) patients (Cave et al., 2017; Rabin et al., 2010). Given the present findings, however, GDE2 dysfunction could well result from defective endocytic trafficking rather than from loss-of-function mutations. Indeed, impaired membrane trafficking is a hallmark of neurodegenerative diseases, including ALS, Parkinson’s and Alzheimer’s disease, and involves Rab GTPase dysfunction, endosomal misrouting and disturbed intracellular transport (Agola et al., 2011; De Vos and Hafezparast, 2017; Kiral et al., 2018; McMillan et al., 2017; Parakh et al., 2018; Schreij et al., 2016; Xu et al., 2018). Thus, disease-associated trafficking defects, even if subtle, could compromise GDE2’s neuroprotective and differentiation-promoting function and thereby contribute to neurodegeneration and possibly other nervous system disorders, an intriguing scenario that merits further study.

## Experimental Procedures

### Cells

SH-SY5Y, N1E-115 and HEK293 cells were grown in Dulbecco’s modified Eagle’s medium (DMEM) supplemented with 10% fetal bovine serum (FBS) at 37°C under 5% CO2. Antibodies used: anti-HA, 3F10 from Roche Diagnostics; β-Actin (AC-15) from Sigma; anti-mCh (16D7) was from Thermo Fisher; anti-EEA1 and LAMP-1 from Abcam; anti-GPC6 antibody LS-C36518 (LifeSpan Bioscience); APC anti-HA Epitope Tag Antibody (Biolegend); EZ-Link™ Sulfo-NHS-Biotin and Streptavidin Agarose Resins were from Pierce; GFP- or mCherry Trap® beads from ChomoTek; Fugene 6 from Invitrogen.

### Plasmids and transfections

Human GDE2 cDNA was subcloned in pcDNA3.1 as described (Matas-Rico et al., 2016). Truncated versions of GDE2 were generated by amplification of full-length GDE2-HA or GDE2-mCherry using reverse primers for the last residues of each truncation. This was followed by a digestion with *BamH*I and *EcoR*V, after which the amplified inserts were cloned into digested and gel-purified pcDNA3.1, and selected by Ampicillin. GDE2 point mutants were generated by site-directed mutagenesis using two complementary oligonucleotides with the desired mutated bases at the center of their sequences. A temperature gradient from 55 to 60 degrees was used during the PCR amplifications. The PCR products were digested with *Dpn*I and transformed into DH5-α competent bacteria and screened for the expected mutated bases.

### Confocal and super-resolution microscopy

Cells cultured on 24 mm, #1,5 coverslips were washed and fixed with 4% PFA, permeabilized with 0.1% Triton X-100 and blocked with 5% BSA for 1 hr. Incubation with primary antibodies was done for 1 hr, followed by incubation with Alexa-conjugated antibodies for 45 min at room temperature. For confocal microscopy, cells were washed with PBS, mounted with Immnuno-MountTM (Thermo Scientific) and visualized on a LEICA TCS-SP5 confocal microscopy (63× objective). Super-resolution imaging was done using an SR-GSD Leica microscope equipped with an oxygen scavenging system, as previously described (Matas-Rico et al., 2016). In short, 15000 frames were taken in TIRF or EPI mode at 10 ms exposure time. Movies were analyzed and corrected using the ImageJ plugin Thunderstorm (http://imagej.nih.gov/ij/), followed by correction with an ImageJ macro using the plugin Image Stabilizer.

### Live-imaging

Live-cell imaging was done on a Leica TCS SP5 confocal microscope equipped with 63x oil immersion lens (numerical aperture 1.4; Leica, Mannheim, Germany). Coverslips were mounted on a metal ring system and exposed to buffer solution (140 mM, NaCl, 5 mM KCl, 2 mM MgCl2, 1 mM CaCl2, 23mM NaHCO3, 10 mM HEPES, 10 mM glucose). N1E-115 cells were selected randomly and images were collected at appropriate time intervals (5-15 sec). GDE2-mCh was visualized by exciting cells at 561 nm, while emission was detected at 610 ± 10 nm.

### GDE2 plasma membrane localization

We performed image analysis for plasma membrane localization of HA-tagged GDE2 constructs by using public domain software IMAGEJ. Shortly, confocal images stained for GDE2-HA were segmented and analysed using Fiji software and a macro that automated the process. First, images were thresholded by the MaxEntropy algorithm to delimit single cells and filtered by Gaussian Blur (radius = 2) and smoothed for segmentation with a median radius of two. Using the Region of Interest manager on Fiji, the background was delimited by using the Li algorithm for thresholding. The cytoplasmic regions were selected by subtracting the plasma membrane thickness (fixed to 0.5 μm, but adjustable from a 0.2-5.0 range) and eroded with a pixel width of one to avoid having empty membranes in segmented cells. Next, the plasma membrane region was obtained by subtracting the background to the cytoplasmic region. Finally, the ratio membrane/cytoplasm was calculated from the median of these regions.

### Western blotting

For Western blotting, cells were washed with cold PBS, lysed in NP-40 buffer supplemented with protease inhibitors and spun down. Protein concentration was measured using a BCA protein assay kit (Pierce) and LDS sample buffer (NuPAGE, Invitrogen) was added to the lysate or directly to the medium. Equal amounts were loaded on SDS-PAGE pre-cast gradient gels (4–12% Nu-Page Bis-Tris, Invitrogen), followed by transfer to nitrocellulose membrane. Non-specific protein binding was blocked by 5% skimmed milk in TBST; primary antibodies were incubated overnight at 4°C in TBST with 2.5% skimmed milk. Secondary antibodies conjugated to horseradish peroxidase (DAKO, Glostrup, Denmark) were incubated for 1 hr at room temperature; proteins were detected using ECL Western blot reagent.

### Biotin labeling

For quantitation of GDE2 internalization and recycling, we used a biotin labeling assay. GDE2-mCh-expressing N1E-115 cells were serum starved for 1hr., transferred to ice, washed in ice-cold PBS, and surface labeled at 4°C with 0.2 mg/ml NHS-SS-biotin (Pierce). For GDE2 internalization, cells were exposed to serum-free medium at 37°C for the indicated time periods. Cells were transferred to ice and washed with PBS, remaining surface biotin was reduced with sodium 2-mercaptoethane sulfonate (MesNa), and the reaction was quenched with iodoacetamide (IAA) prior to cell lysis. For recycling assays, cells were labeled with biotin as above, and incubated in serum-free medium at 37°C for 30 min to allow internalization of GDE2. Cells were returned to ice, washed with PBS, and biotin was reduced using MesNa. Recycling of the internal GDE2 pool was induced by a temperature shift to 37°C for 0–30 min. Cells were returned to ice, washed with PBS and surface biotin was reduced by MesNa. MesNa was quenched by IAA and the cells were lysed. Biotin-labeled GDE2 was detected using Streptavidin beads and anti-mCh antibody.

### Immunoprecipitation

For co-immunoprecipitation of GDE2 and Rabs, HEK293Tcells were plated on plastic dishes of 10 cm diameter and transient co-transfected with GDE2-mCh or –GFP and Rab4-GFP, Rab5-mCh, Rab7-GFP or Rab11-mCh. After 24 hrs cells were lysed using RIPA buffer. Protein concentration was determined using Protein BCA protein assay kit (Pierce). Immunoprecipitation was carried out incubating 500 μg - 1 mg cytoplasmic extracts with GFP- or mCherry Trap® beads(ChomoTek) at 4°C for 1hr. Beads were washed three times and eluted by boiling in SDS sample buffer for 10 min. at 95°C. Supernatants were applied onto an SDS gel and subjected to immunoblot analysis

### Inducible GDE2 expression

SH-SY5Y cells with inducible expression of GDE2 constructs were generated using the Retro-X™ Tet-On® Advanced Inducible Expression System (ClonTech), as described (Matas-Rico et al., 2016). After retroviral transduction, the cells were placed under selection with G418 (800 mg/ml) supplemented with puromycin (1 μg/ml) for 10 days. GDE2 induction (in the presence of 1 μg/ml doxycycline) was verified by Western blot and confocal microscopy. Transient transfection was performed with Fugene 6 reagent (Invitrogen) according to the manufacturer’s instructions.

### Flow Cytometry

For GPC6 and GDE2-HA surface expression analysis, cells were grown in complete medium with 10% FCS with or without doxycycline. Cells were trypsinized into single-cell suspensions and then 8×10^5^ cells were incubated with 5 μl of anti-GPC6 antibody LS-C36518 (LifeSpan Bioscience) and in 4 μl of APC anti-HA antibody (Biolegend). Bound GPC6 antibody were detected by incubating with a 1:200 dilution of goat anti-mouse Alexa-488 secondary antibody in 2% BSA for 45 min on ice. Fluorescence measurements were performed using BD LSRFORTESSA and using Flow Jo software.

### Induction of neuronal differentiation markers

Total RNA was extracted using the GeneJET purification kit (Fermentas). cDNA was synthesized by reverse transcription from 5 μg RNA using First Strand cDNA Syntesis Kit (Thermo Scientific). RTqPCR was performed on a 7500 Fast System (Applied Biosystems) as follows: 95°C for 2 min, 95 °C for 0 min., 40 cycles at 95°C for 15 sec. followed by 60°C for 1 min. for annealing and extension. The final reaction mixtures (20 μl) consisted of diluted cDNA, 16SYBR Green Supermix (Applied Biosystems), 200 nM forward primer and 200 nM reverse primer. Reactions were performed in 96-well plates in triplo. The primers used are listed below. As a negative control, the cDNA was replaced by milliQ water. Cyclophilin was used as reference gene. The normalized expression (NE) data were calculated by the equation NE= 2(Ct target – Ct reference).

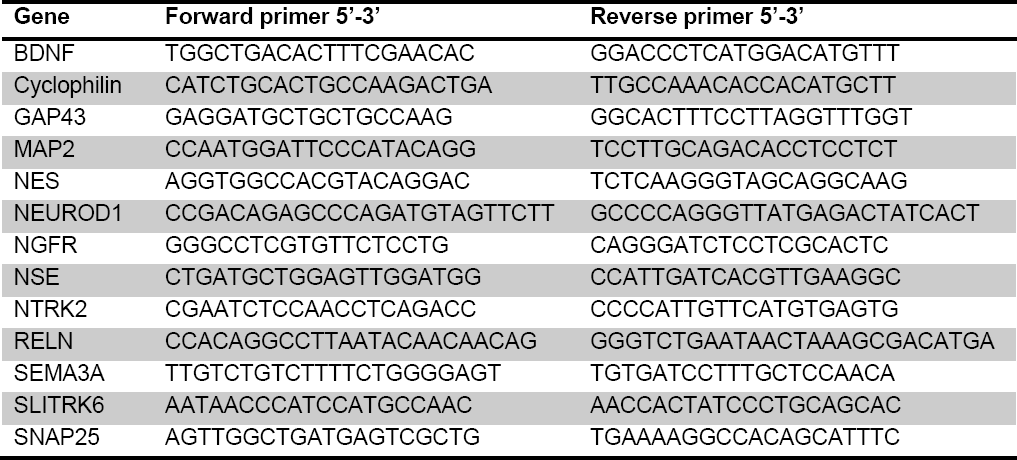

## Supplemental Information

Supplemental Information includes three figures and one movie.

## Acknowledgments

This study was funded by the Dutch Cancer Society (KWF; project 2017-10215).

## Author Contributions

F.S.-P., M.v.V., B.v.d.B., D.L.-P., and E.M.-R. conceived and performed experiments. A.P., W.H.M., and E.M.-R. supervised the work. W.H.M wrote the manuscript with feedback from F.S.-P., A.P., and E.M.-R.

## Supplemental Information

**Movie S1. Live imaging of GDE2-containing vesicle movements in SH-SY5Y cells.**

**Figure S1.**
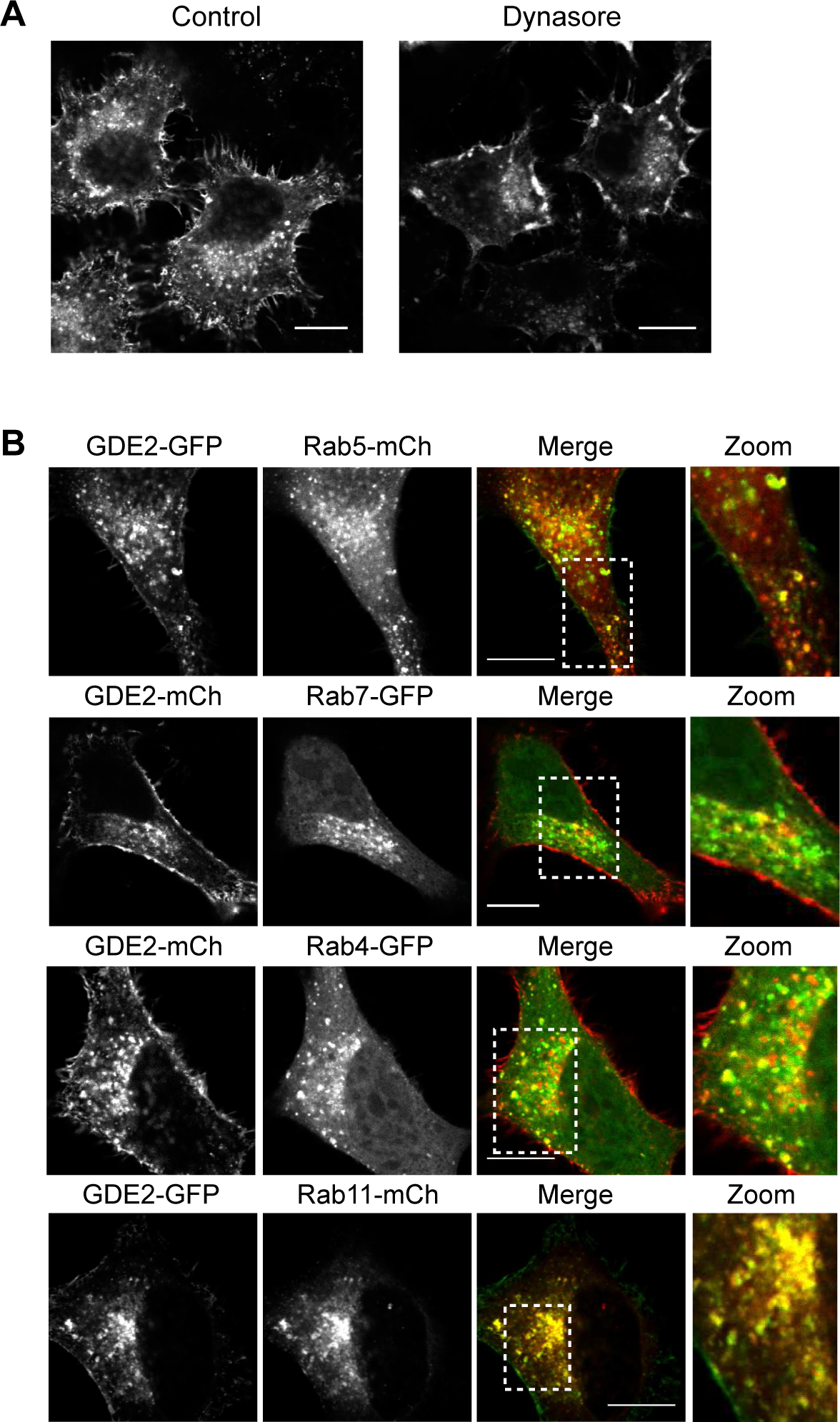
GDE2 subcellular localization in N11E-115 cells. **A.** GDE2 accumulates at the cell surface with loss of GDE2-positive vesicles in N1E-115 cells treated with the dynamin inhibitor Dynasore (80 µM). Bar, 10 µm. **B.** GDE2 co-localization with the indicated Rab GTPases in N1E-115 cells. Bar, 10 µm. See Fig. 1D and text for details.

**Figure S2.**
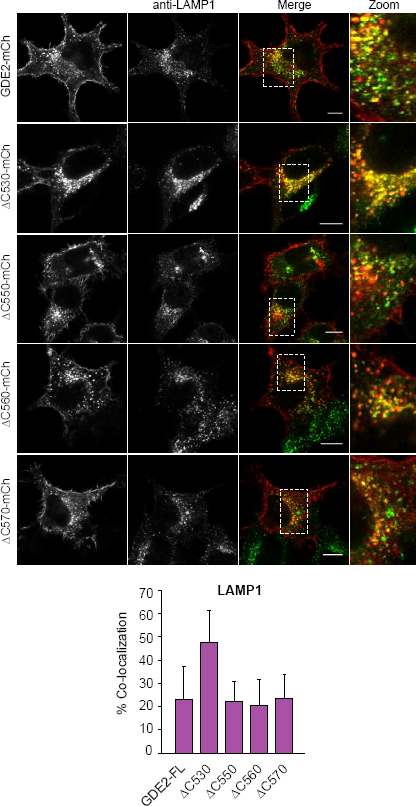
Lysosomal localization of GDE2 and its truncation mutants. N1E-115 cells expressing the indicated GDE2-mCh constructs were immunostained for LAMP1, using LAMP1-specific antibody. Bar, 10 μm. Lower panel, quantification of GDE2-LAMP1 co-localization. Data represent the median ± SEM (error bars) of co-localization.

**Figure S3.**
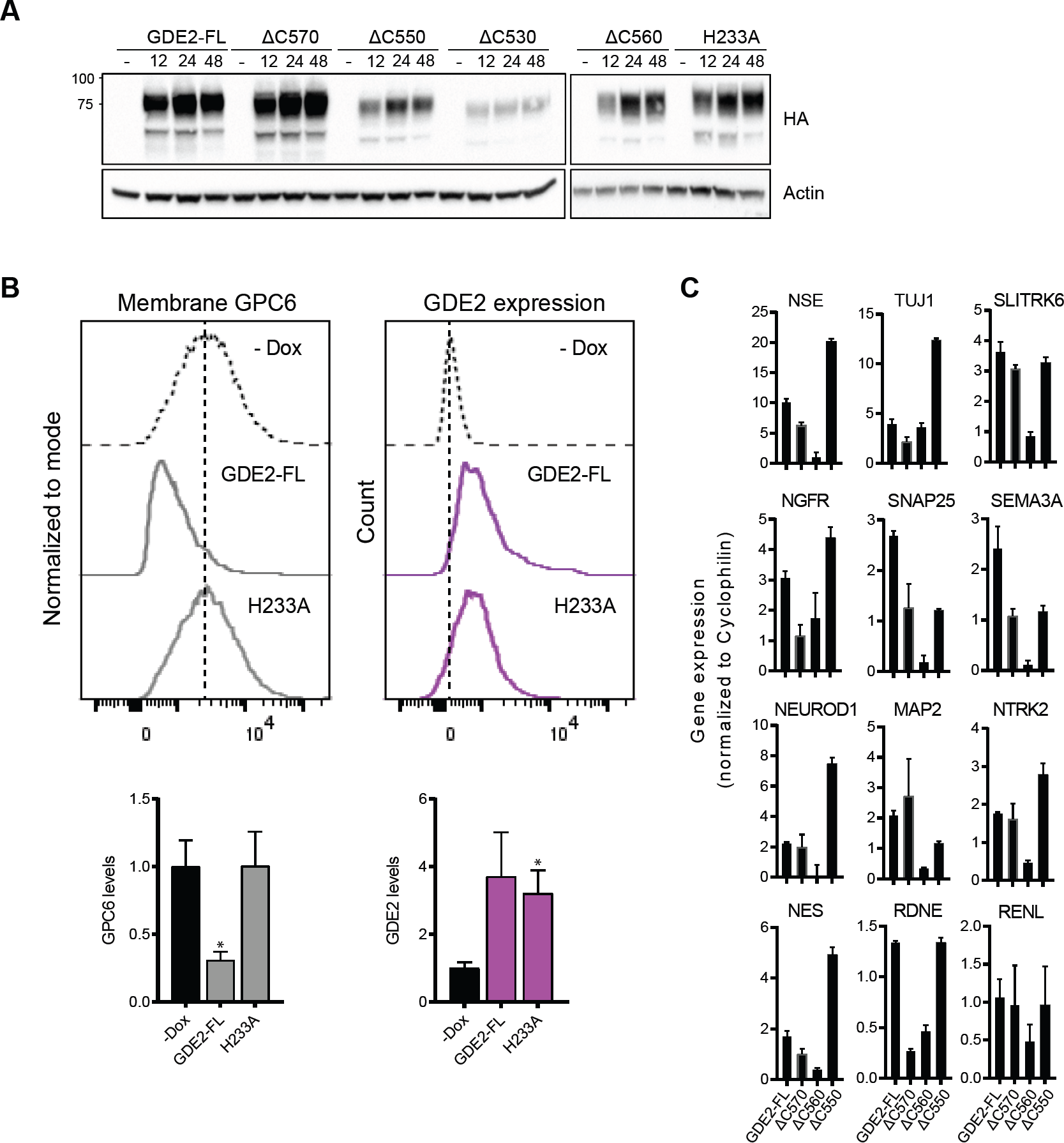
Expression and localization of GDE2 and truncation mutants in Dox-treated SH-SY5Ycells. **A.** Western blots showing expression of the indicated GDE2-HA constructs after 24 and 48 hrs of Dox treatment. Actin was used as loading control. See also Fig. 3A. **B.** Control experiment showing that GDE2(H233A) at the cell surface is inactive towards GPC6. Cell surface expressions were determined by FACS. Data show the median ± S.E. (error bars) from triplicate measurements taken from two independent experiments. *p<0.05, paired t test. See also Fig. 3C. **C.** Induction of neuronal differentiation genes upon Doxycycline induction (48 hrs) of the indicated GDE2 constructs, as determined by qPCR. Data represent the average value of triplicate measures ± SEM (error bars) analyzed by paired t test. See also Fig. 3D.

